# A Universal Phase Transition in Plankton Trait Dynamics

**DOI:** 10.1101/679209

**Authors:** Jenny Held, Tom Lorimer, Ruedi Stoop, Francesco Pomati, Carlo Albert

## Abstract

Key ecological traits, like cell size, often follow scale-free or self-similar distributions. This indicates that these systems might operate near a critical (i.e. second-order) phase transition where macroscopic system behaviour is largely decoupled from microscopic system details, allowing an extremely simple, yet accurate and robust mathematical system characterisation. However, how trait-distribution scaling results from a critical transition has not yet been explicitly demonstrated. Here, we demonstrate that a generic class of cell growth and division models exhibits a critical transition from a growth-dominated to a division-dominated phase. We find experimental evidence for this transition, both in the population dynamics and in the moment scaling of chlorophyll distributions, for prokaryotic and eukaryotic phytoplankton growth under different light intensities. Our approach offers testable predictions of the response of unicellular trait-distributions to perturbations.

## 1 Introduction

Scale-freeness, manifesting as power law distributions over many orders of magnitude, is routinely observed in complex natural and artificial systems (Rinaldo et al., 2002; Banavar et al., 2007; Chialvo, 2010; Lux and Marchesi, 1999; Faloutsos et al., 1999). From the theory of interacting particles we know that scale-freeness is often associated with critical phenomena. Close to a critical point, a system’s behaviour is accurately described by few system-level observables that follow simple universal scaling laws with respect to the few relevant variables that measure the deviation from the critical point. These scaling laws are valid for whole classes of systems, even though the systems may differ in many details, which greatly simplifies mathematical modelling and prediction. Natural systems might operate near critical points because diverging fluctuation correlation lengths cause high susceptibility to external stimuli, which can enhance adaptive responses in complex systems (Kern and Stoop, 2003; Stoop and Gomez, 2016; Kinouchi and Copelli, 2006). Theories have also been put forward to explain how complex systems might arrive at criticality through a self-organising process (e.g. Bak et al., 1987). In the field of ecology, this general idea has been used to explain, for instance, scale-free species-abundance distributions (Solé et al., 2002). While the presence of a critical point is therefore of great theoretical and practical interest, direct demonstrations that observed scaling laws in ecology result from a critical point are generally missing. To this end, we demonstrate here that a generic class of ecological models of unicellular organisms exhibits a critical point, and provide experimental evidence for this critical point in cellular level trait distributions.

We will focus on aquatic unicellular organisms, which are of fundamental importance for global nutrient cycles. These organisms can be characterised by traits such as size, biomass, or pigment content, that are often distributed in a scale-free manner across individuals. For instance, size distributions have been observed to be scale-free over almost 3 orders of magnitude (Rinaldo et al., 2002; Marañón, 2015; Cavender-Bares et al., 2001). Moreover, laboratory experiments have shown that under ideal growth conditions, size distributions of many different unicellular aquatic species are self-similar, that is, they collapse onto a universal function under re-scaling (Giometto et al., 2013). Here we show that also the distributions of cellular chlorophyll content under different light intensities are self-similar under certain environmental conditions. Furthermore, we present a model class that incorporates the most relevant processes of unicellular population and trait dynamics: trait increase at the cellular level (which we shall henceforth refer to as ‘growth’), cell division, and cell death. Our model class incorporates three time scales, and two trait scales arising from limiting processes that prevent cells from having trait values that are too large or too small. Within this class, we find a critical point marking a transition from a growth-dominated phase (GDP) to a division-dominated phase (DDP). In the vicinity of the critical point, the mean trait value of the population obeys power laws with respect to both the natural time scale of trait dynamics and the limiting trait scales. These scaling laws are largely independent of the details of the growth, division, and death processes. They originate from the fact that the underlying universal power law characterizing the trait distributions has an exponent that jumps from −1 to −2 at the phase transition. These power laws can be masked by the limiting processes and hence may not always be visible. We show, however, that even in these cases, it is possible to find these power laws in the scaling of the higher moments of trait distributions across different populations (for example, corresponding to different species or different environmental conditions).

After presenting our class of models in Section 2.1, we describe our experimental data in Section 2.2. In Section 3.1, we demonstrate theoretically the existence of a phase transition in our model class. We then present experimental evidence for this phase transition in size- and chlorophyll-distributions in Section 3.2. In Section 4 we discuss some of the wider implications of our findings and suggest experiments to test its further predictions.

## 2 Methods

### 2.1 Growth-death-division model

We model a population of cells, where the state of each cell is completely described by a single positive continuous trait value *x*. Here, we consider ‘traits’ like biomass or chlorophyll content that vary continuously at cellular level (except upon cell division). The rates associated with the processes considered here (trait increase or ‘growth’, cell division, cell death) are assumed to be functions of *x* that will of course depend on the species and on the environmental conditions. We denote the *growth* rate of an individual cell per unit of time as *g*(*x*) (i.e. *dx/dt* = *g*(*x*)). The *cell division* and *cell death* rates, i.e. the instantaneous division and death probabilities, are denoted as *b*(*x*) and *d*(*x*), respectively. Upon division, the cell splits into two daughter cells, each with trait value *x/*2. Such a division process makes this class of models inherently *non-local* in trait space. Under the assumption of a large number of cells, we may neglect cell-number fluctuations, and describe the system with a deterministic trait- and time-dependent cell number density, *ψ*(*x, t*), which satisfies the linear, non-local PDE (Hall and Wake, 1989, 1990),

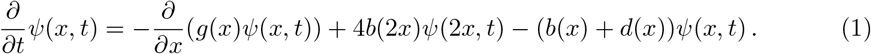

The rates *g*(*x*), *b*(*x*), and *d*(*x*) are required to be positive (functional forms will be discussed later). Somatic growth, described by the term 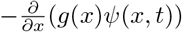, induces a drift of cells to higher trait values. The non-local term 4*b*(2*x*)*ψ*(2*x, t*) describes cells appearing at *x* due to divisions at 2*x*. The factor 4 is a product of a factor 2 (because at division, 2 new cells are created) and the Jacobian resulting from the transformation *x* ↦ 2*x*. The last term on the r.h.s. of Equation (1) encodes loss of cells at trait values *x* due to division and death, respectively.

Consider first the base model where the rates have the functional forms *g*(*x*) = *ω*_1_*x*, *b*(*x*) = *ω*_2_, and *d*(*x*) = *ω*_3_, respectively. While *ω*_1_ quantifies the exponential growth at cellular level, *ω*_2_ and *ω*_3_ quantify the population growth. This basic model does not contain trait scales, since all parameters (*ω*_1_, *ω*_2_, and *ω*_3_) are *frequencies*. The population will grow exponentially without bound (considering only cases where *ω*_3_ < *ω*_2_), yet may still reach an equilibrium trait-distribution. Because of the absence of trait scales, however, the only possible equilibrium trait-distribution is a power law distribution (the only scale-free distribution). Since pure power laws cannot be normalised, however, there cannot be an equilibrium trait-distribution for this model.

In reality, cells have a limited range of accessible trait values, which resolves the normalization issue. To implement these limits, our model class discourages small trait values by increasing the growth rate at small trait values, and discourages large trait values by increasing the division rate there. The death rate is assumed to be trait-independent. The precise – mathematical or biological – implementation of these two limiting processes does not matter for the universal scaling results that we will present here (a precise description of the model class is given in Section S1 of the Supplementary Material). We only require these limiting processes to be strong enough for an equilibrium trait-distribution with finite moments to exist. The limiting processes introduce a lower trait scale, *u*, and an upper trait scale, *v*. The values of these trait scales will depend on the particular trait and the species we are interested in, and on the environmental conditions (e.g. the light intensity, if the trait of interest is chlorophyll content). In general, more than two trait scales might be needed to describe realistic growth and division processes. However, it turns out that the scaling phenomena we observe are well described by at most two scales.

After a sufficiently long time (compared to the trait relaxation time scale of the model |*ω*_1_ − *ω*_2_|^−1^; see Section S1), the trait-distribution will reach an equilibrium while the cell number continues to increase at a steady exponential rate. The state of the system may thus be described by equation

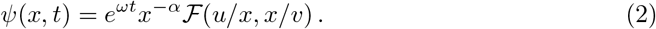

In this ansatz, *α* is a function only of *ω*_1_ and *ω*_2_ (see Section S1), whereas 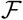 is a general cut-off function that will depend on the details of the limiting processes. For small arguments, 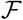 returns values of approximately unity, otherwise 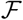 quickly approaches zero. Thus, the time invariant trait-distribution will be a power law in the range *u* ≪ *x* ≪ *v*. The effective population growth rate *ω* is a function of all the model parameters (*ω*_1_, *ω*_2_, *ω*_3_, *u*, and *v*). The dependence of *α* and *ω* on the parameters of our model class will be investigated in Section 3.1.

In empirical data, exponent *α* may not always be directly observable, as the power law *x*^−*α*^ in Equation (2) may be masked by the cut-offs. In such cases, we can still observe this exponent indirectly. If a whole family of trait-distributions shares the same 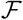 but with different values of *u* and *v*, then *α* can indirectly be obtained through *moment scaling*, by observing how the higher moments of the trait distribution scale with the first moment, the mean trait (see Sections S1 and S2.2). Such a family of distributions may for instance be obtained by taking measurements of different species, or of single species under different environmental conditions (as we will do in Section 3.2). Depending on the value of *α* and the variation of the trait scales in the family, a *data collapse* of the observed trait distributions onto a single universal curve 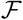 may be observed upon re-scaling with the distributions’ mean trait values (see also Section 3.2).

In order to corroborate our theoretical results and experimental evidence of the phase transition, we also ran simulations with a member of our model class. We generalised the *Gillespie algorithm* (Gillespie, 1976) to systems with variable rates (described in detail in S3.2) in order to qualitatively reproduce the experimental results (see Section S3.1).

### 2.2 Phytoplankton chlorophyll content under different light intensities

We use chlorophyll-a measurements in phytoplankton monocultures as experimental support for our theoretical findings (Section 3). Our chlorophyll measurements were derived from experimental data described in detail in Fontana et al. (2019), which we summarise briefly here. *Kirchneriella subcapitata* (a eukaryotic green alga) and *Microcystis aeruginosa* (a prokaryotic cyanobacterium) were grown in monocultures under several different, but constant, light intensities and a constant temperature of 20°C. All cultures were subjected to the same light-dark cycle, with 14 hours light and 10 hours darkness, and measurements were taken during the light period. All cultures received the same initial nutrient concentration, and nutrients were not replenished. The cultures were stirred regularly to avoid stratification and self-shading. Of each species there were six different directly illuminated light intensity treatments, with two replicates each. Additionally, for each replicate (species and light intensity), there was an additional monoculture of the other species in a separate container, that was shaded by the first (no mixing). The mixed cultures described in Fontana et al. (2019) are not considered here.

The data were collected using a scanning flow cytometer as described in Fontana et al. (2019), from which we only analyse live cells that were identified through clustering and a trained random forest classification algorithm (Thomas et al., 2018). We look here at the *total fluorescence responses* (that is, the integral over the fluorescence pulse) of the cells in the red spectrum (677–700 nm). This channel captures mainly the fluorescence of the pigment chlorophyll-a, responsible for photosynthesis and thus relevant for the energy acquisition of the cell.

From previous studies it is known that the fluorescence and the chlorophyll content of phytoplankton cells are linked by a scaling relationship of the form ⟨*f*⟩ ~ ⟨*x*⟩^ϵ^, *ϵ* < 1, across cultures of the same species subjected to different light intensities, where ⟨*x*⟩ is the mean chlorophyll content and ⟨*f*⟩ the mean chlorophyll fluorescence of a culture (Álvarez et al., 2017). The scaling exponent *ϵ* < 1 is thought to be the result of the pigment distribution inside the cells, and the fact that pigments in the body of the cell receive less light and contribute less to fluorescence than pigments close to the cell wall (‘package effect’), resulting in sub-linear scaling (Álvarez et al., 2017). Such a power law transformation does not change the scaling relationships we discuss here. Nevertheless, all fluorescence data presented here have been power law transformed, in order to present chlorophyll estimates rather than fluorescence measurements. The value of *ϵ* is not exactly known for the instrument and species we consider here, so with the only constraint *ϵ* < 1, an exponent *ϵ* = 0.5 was chosen. In these data, the power law in Equation (2) cannot be observed directly, but only through moment scaling (Sections 2.1, S1, S2.2), as will be shown in Section 3.2. In order to observe this moment scaling clearly, we remove outliers from the data. A detailed description of this process and the effect it has on the observed scaling relationships is given in Section S2.1, and the biological origins of these outliers are discussed in Section 4.

## 3 Results

### 3.1 Theoretical result: second order phase transition in trait dynamics

In this section, we investigate the dependence of *α* and *ω* in Equation (2) on the parameters of our model class. To this end, we introduce the dimensionless parameters *h* = *u/v* and *τ* = *ω*_2_*/ω*_1_ − 1, which are relevant for the trait dynamics. The former defines the lower trait scale relative to the reference scale *v*. The latter defines the fundamental time scale of the trait dynamics, relative to the reference time scale 1/*ω*_1_. The *critical point* is at *τ* = 0 and *h* = 0. At this point, trait-distributions follow power laws at trait values well below *v*, and the return to the equilibrium trait-distribution after a perturbation also follows as power law in time, rather than an exponential decay (a phenomenon known as *critical slowing down*, see Section S1). This critical point marks a transition from a growth-dominated phase (GDP), where *ω*_1_ > *ω*_2_, to a division-dominated phase (DDP), where *ω*_2_ > *ω*_1_. At the transition point, the population growth rate *ω* in Equation (2) changes from *ω*_1_ − *ω*_3_ to *ω*_2_ − *ω*_3_, both up to corrections of order *h* (see Section S1). In Equation (2), the exponent *α* is a smooth function of *τ*, except at the critical point, where the value of *α* jumps from 1 (GDP) to 2 (DDP). This jump is responsible for a drastic change in the scaling of the mean trait ⟨*x*⟩ : in the GDP, cells grow faster than they divide, for most trait values, so that ⟨*x*⟩ scales as the upper limiting trait scale *v*. In the DDP, likewise, ⟨*x*⟩ scales as the lower limiting trait scale *u* (see Section S1). This means that, at the transition, the scaling of ⟨*x*⟩ with respect to the limiting trait scales changes from ⟨*x*⟩ ~ *h*^1^ (GDP) to ⟨*x*⟩ ~ *h*^0^ (DDP) (in units where *v* = 1).

More generally, in the *scaling regime* near the critical point, where both *τ* and *h* are small, the behaviour of the system can be expressed through *scaling laws* in terms of *τ* and *h* (see Section S1). For *h* ≪ |*τ*| ≪ 1, and in units where *v* = 1 and *ω*_1_ = 1, they can be captured with the scaling ansatz (see Equation S2)

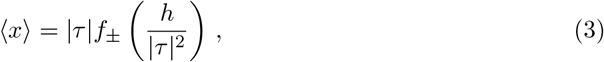

where the functions *f*_±_, valid for positive and negative *τ*, respectively, are analytic functions, for small arguments, and both *f*_+_(*y*)*/y* as well as *f*_−_(*y*) converge to positive constants when *y* → 0. From Equation (3) we derive two scaling laws, respectively, for the mean ⟨*x*⟩ and its *susceptibility* to changes of external conditions (measured in terms of *h*),

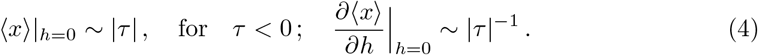

In addition, the time it takes to return to an equilibrium trait distribution after a perturbation diverges according to the power law (see Section S1)

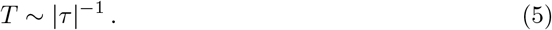

These laws are *universal*: they do not depend on the details of the growth and division processes within our model class (see Section S1). We thus expect to find them in many trait-distributions that are governed by such processes. Potential implications of these scaling laws are discussed in Section 4.

In physics, the kind of transition we describe here is known as a *second order phase transition*, an effect that is classically exemplified by the ferromagnetic-paramagnetic transition at the Curie temperature (see Box 1). The similarity between our biological phase transition and the magnetic phase transition is demonstrated by comparing the behaviour of the order parameters of both transitions near the critical point (see Figure 1). As discussed in Section 2.1, we can observe the jump in the exponent *α* responsible for the phase transition through moment scaling. Near the critical point, the higher moments ⟨*x*^*k*^⟩ (*k* = 2, 3, …) scale with the first moment as ⟨*x*^*k*^⟩ ~ ⟨*x*⟩^*λk*^ across a family of distributions with the same scaling function 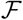 (see Sections S1 and S2.2). In the GDP, as all moments are dominated by cells of large trait values and scale like powers of *v*, we find a trivial moment scaling with *λ* = 1. In the DDP, the mean scales with *u*, but the higher moments, dominated by cells of large trait values, scale like *v*. As *u* and *v* will generally scale differently across the family, we will have a moment scaling with *λ* ≠ 1. In the next section, we give two examples of families of distributions (consisting of different species in one case and different environmental conditions in the other) where the exponent *α* can be observed via moment scaling. In the latter, we even observe the phase transition itself.

**Figure 1:**
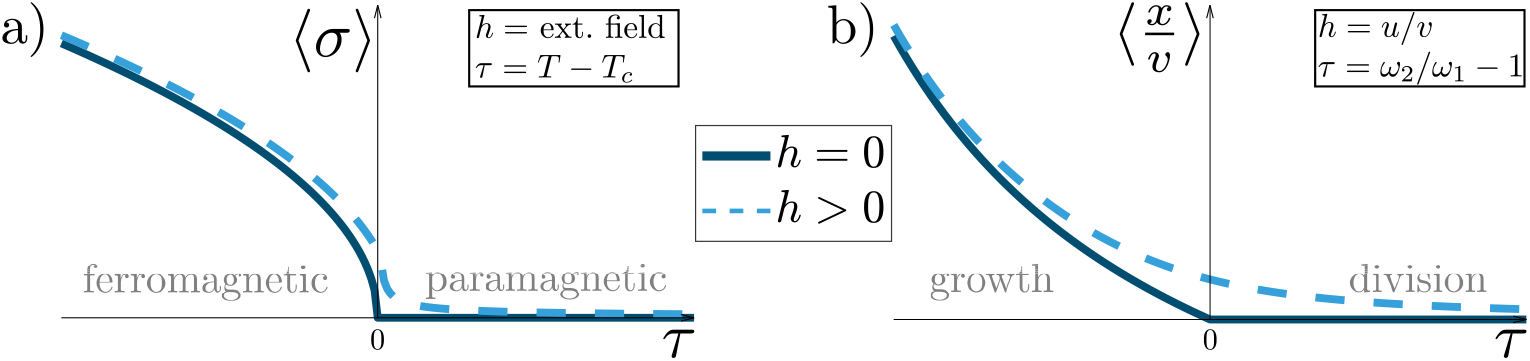
Behaviour of the order parameters in two second order phase transitions. In magnets (a), in the absence of an external field *h*, the magnetization ⟨*σ*⟩ (*order parameter*) is close to zero in the paramagnetic phase, but non-zero in the ferromagnetic phase. In our biological system (b), the normalised mean trait ⟨*x/v*⟩ is the order parameter and is close to zero in the division dominated phase, but non-zero in the growth dominated phase.

##### Box 1: Analogy to the phase transition in magnets.

A simple model for a ferromagnetic material is a lattice of binary units that can adopt either an “up” or a “down” magnetic *spin*. Each unit seeks to align its spin with its neighbours (minimising energy), but thermal fluctuations counteract this natural tendency. If the temperature *T* is above a certain critical temperature *T*_*c*_, thermal noise overpowers the spin alignment and the net internal magnetic field ⟨*σ*⟩ (the *order parameter*) vanishes: the *paramagnetic* phase. Below the critical temperature, patches of spins spontaneously align, and the system develops a net magnetic field: the *ferro-magnetic* phase. In both phases, a small *externally applied* magnetic field will induce a net magnetisation, softening the distinction and transition between these two phases (see Figure 1a).

In our model, the GDP corresponds to the ferromagnetic phase, and the DDP to the paramagnetic phase (see Figure 1b). Our model parameter *τ* corresponds to the temperature, *h* to the externally applied magnetic field, and 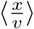 to the net induced magnetic field. Increasing cell division (i.e. increasing *τ*) disturbs growth, like increasing temperature disturbs spin alignment. Increasing *h* pushes the cells to larger trait values relative to *v*, like an external magnetic field pushes the spins to align. The order parameter 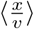 measures how successfully the whole population has grown, like the net magnetisation measures how many spins have aligned.

While the behaviour of the order parameters of these two systems look qualitatively similar (Figure 1), the *critical exponents* measuring how exactly they scale with *τ* are clearly differen In the spin system, in the ferromagnetic phase the order parameter increases with 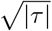, whereas in the population model, in the corresponding growth-dominated phase, it increases linearly with |*τ*|, for small |*τ*|, according to the l.h.s. of Equation (4).

### 3.2 Experimental evidence for the phase transition

#### Size-scaling of unicellular aquatic species in vitro

The equilibrium size distributions, *ψ*_i_(*x*), of various unicellular aquatic species (indexed by *i*) have been found to be, to a good approximation, self-similar upon re-scaling with the species’ mean sizes ⟨*x*⟩_*i*_ (Giometto et al., 2013). That is, for the trait *x* being ‘cell size’, the size distribution of species *i* can approximately be described by 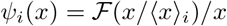, which is consistent with Equation (2) when *α* is close to but smaller than 1 (because then the parameters *v*_*i*_ scale like ⟨*x*⟩_*i*_) and the parameters *u*_*i*_ do not vary significantly. The resulting moment scaling 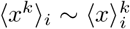 was experimentally confirmed up to *k* = 4. These observations are consistent with our model under the assumption that the populations were in their GDP (see Sections S1 and S2.2).

#### Chlorophyll-fluorescence under different light intensities

Using flow cytometry data from Fontana et al. (2019) (cf. Section 2.2), we analysed single-cell absolute chlorophyll content in phytoplankton monocultures exposed to different light intensities.

In all cultures, we observe initially fast population growth of exponential form, with roughly 2 days doubling time (Figure 2a). At the same time, all positive moments of the chlorophyll content distributions across the light intensities scale trivially as powers of a single trait scale, for instance the mean chlorophyll content ⟨*x*⟩, for both cell types: we find ⟨*x*^*k*^⟩ ~ ⟨*x*⟩^*k*^, for *k* = 2, 3, …, 5 (Figure 3a; up to 5 moments can be reliably estimated after removal of outliers, c.f. Section S2.1). The data also show an approximate data collapse (Figure 4c; c.f. Section 2.1).

**Figure 2:**
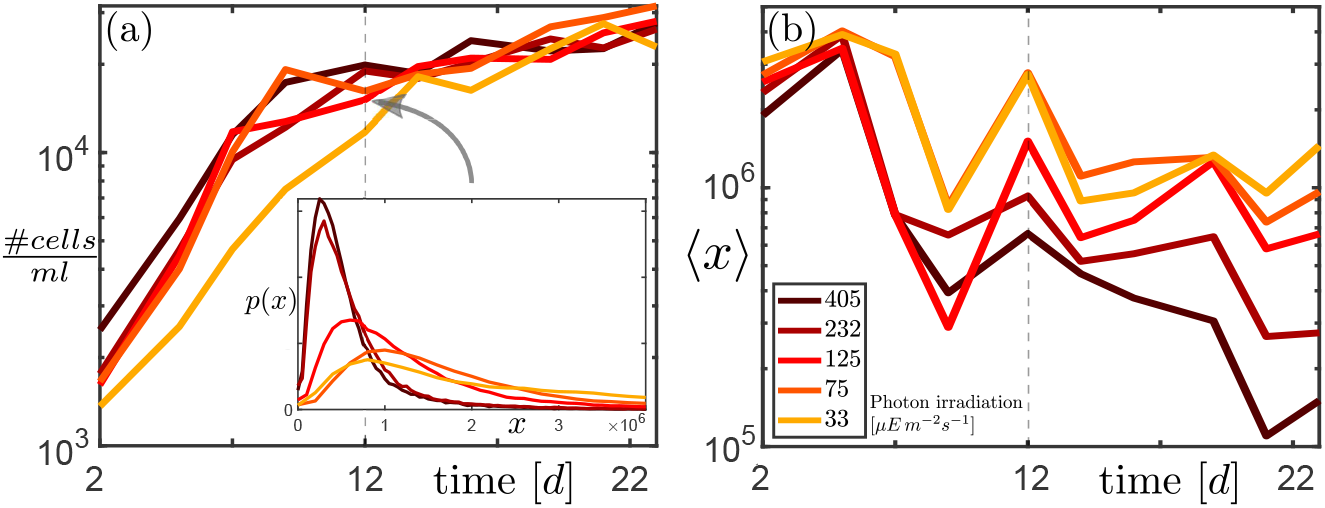
(a) Cell density in green algae cultures exhibiting a transition from fast to slow population growth. *Inset:* Distributions *p*(*x*) of absolute cell chlorophyll content *x* shortly before transition. (b) Mean absolute cell chlorophyll development. Colors indicate the light irradiation intensity. The dotted grey line marks roughly the transition point. Plots from other green algae and cyanobacteria cultures are shown in Section S2.1.

**Figure 3:**
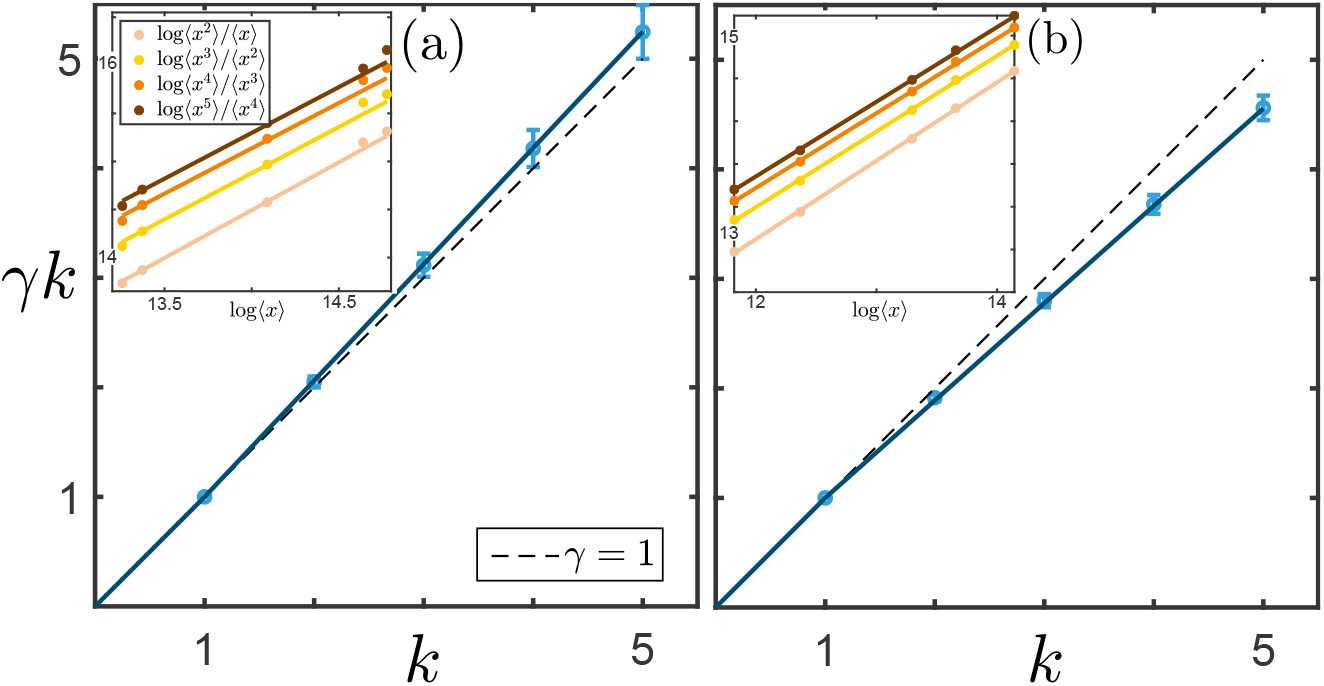
*γk* = log ⟨*x*^k^⟩/ log ⟨*x*⟩ : scaling between first and higher moments. (a) Before the transition, trivial scaling (⟨*x*^*k*^⟩ ~ ⟨x⟩^*k*^ is observed, (b) after the transition, higher moments scale differently (⟨*x*^*k*^⟩ ~ ⟨*x*⟩^*γk*^, with *γ* < 1). Two points in time are shown (a: day 12, b: day 22), but the moment scaling stays similar to (a) before the transition, and similar to (b) after the transition. Plots from other green algae and cyanobacteria cultures are shown in Section S2.1. The error bars indicate the 95% confidence intervals for the exponents *γk*, that were obtained by fitting a linear regression model to the log-transformed moments. Before the fit, outliers were removed (see Sections 3.2 and S2.1). *Insets:* Moment ratios, where points along each line correspond to the different light intensities. The slopes of the straight line fits determine the exponents *γk*.

**Figure 4:**
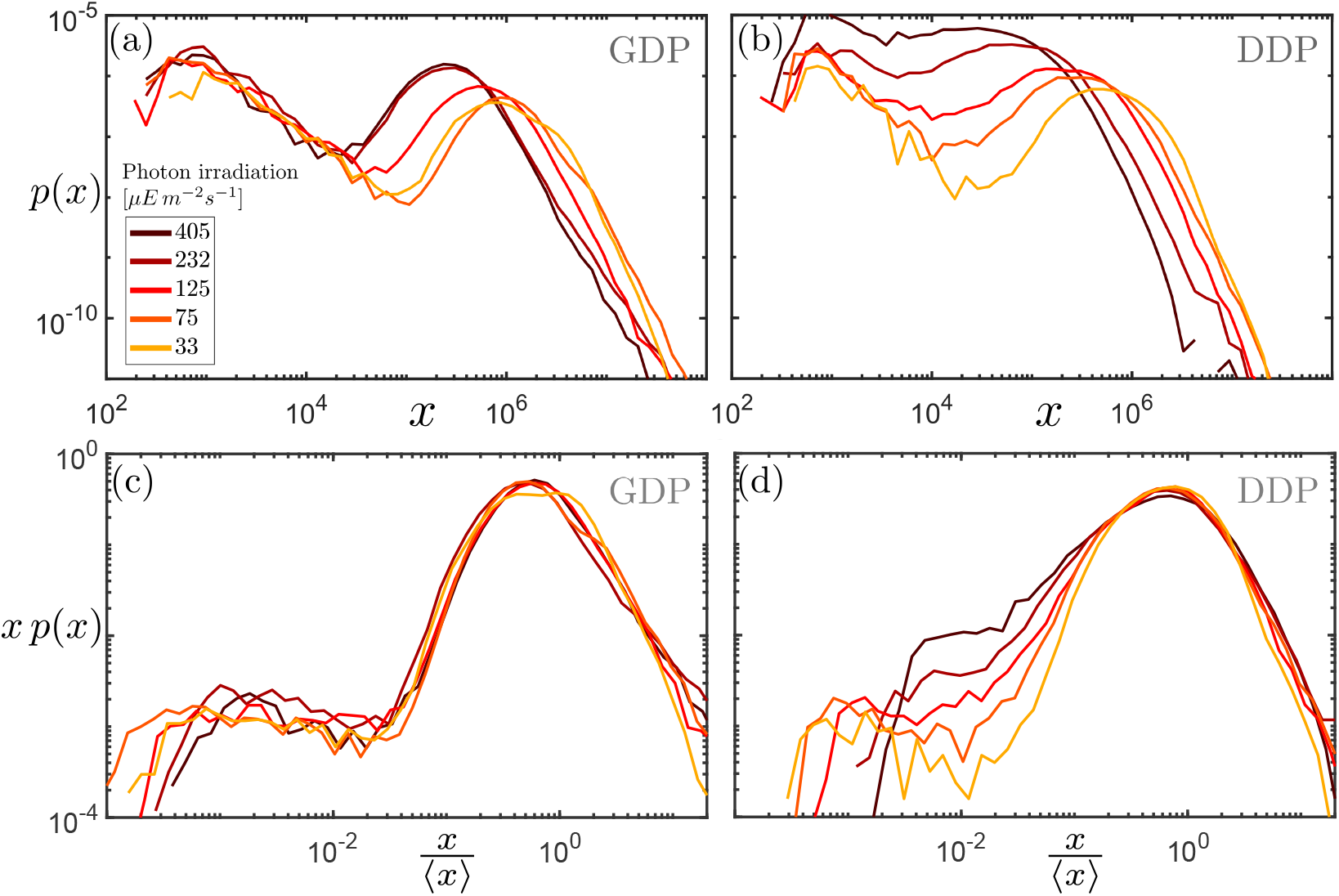
Distributions *p*(*x*) of absolute chlorophyll content *x* of cells in the green algae cultures before (a,b) and after rescaling (c,d), at two dates, one before and one after the transition (day 12 (a,c) and day 22 (b,d), respectively). A data collapse is obtained before the transition only, whereas after the transition the distributions no longer collapse well onto a single curve.

After the transition, the mean chlorophyll content of cells decreases strongly, reaching approximately an order of magnitude less than the initial amount by the end of the experiment (Figure 2b). At the same time, the overall population growth slows down significantly, showing a doubling time of roughly an order of magnitude less than in the initial phase (Figure 2a). Simultaneously, the higher moments *k* = 2, 3*, … ,* 5 still scale as powers of the first moment ⟨*x*⟩, but now as ⟨*x*^*k*^⟩ ~ ⟨*x*⟩^*γk*^, with *γ* < 1 (Figure 3b). Consequently, there is no longer a data collapse (Figure 4d).

Applying our model (Section 2.1) to these data, the observed moment scaling is consistent with the underlying power law exponent *α* being smaller but staying close to 1 in the initial phase, and being larger but staying close to 2 in the later phase (see Section S2.2). In both phases, we also observe the previously described lower cellular chlorophyll content in higher light intensity settings (our Figure 2b; MacIntyre et al. 2002; Álvarez et al. 2017).

### 4 Discussion

The experimental data in Section 3.2 provide evidence for our theoretically predicted two phases (GDP and DDP), the transition between these two phases, and most importantly, the predicted scaling behaviour, *in both of the phases*, and throughout the experiment. The evidence for the GDP and DDP, and the transition, can be seen in the shift from a higher (GDP) to a lower (DDP) population growth rate and mean chlorophyll content nearly simultaneously (Figure 2). This connection between the population growth rates and the chlorophyll content is a quintessential feature of the two phases (see Section S2.2). In the GDP, the population growth rate *ω* is dominated by the growth rate *ω*_1_, and the mean chlorophyll content ⟨*x*⟩ is dominated by the upper cut-off *v*. In the DDP, *ω* is dominated by the division rate *ω*_2_, because *ω*_1_ has dropped below *ω*_2_, and *x* is dominated by the much smaller lower cut-off *u*. In our data, the decrease in the population growth rate after the transition appears to be enhanced by an increased death rate *ω*_3_ (see Section S3.1). The evidence of scaling in both of the phases can be seen in the trivial moment scaling and approximate data collapse before the transition, and non-trivial moment scaling without data collapse after the transition (Figures 3, 4). This is consistent with our theory for both the GDP and DDP respectively, near the critical point of our model class (for derivations, see Section S2.2). This scaling evidence is crucial, as it suggests that this very complex biological system actually remains sufficiently close to the critical point of our phase transition *throughout the experiment*, that its behaviour can be simply and robustly described by our universality class. That is, even though the phase transition itself occurs abruptly, it still dominates the experimental system behaviour for several days both before and after it occurs. In our experimental data, the phase transition might have occurred due to nutrient limitation (e.g. by depletion of carbon), as nutrients were not replenished during the experiment. The critical point can dominate the system behaviour even in cases where the transition does not occur, and the system remains in only one of the phases. In the data of Giometto et al. (2013), for example, where nutrients were not limiting, the data collapse and trivial moment scaling in the size distributions suggest that their cell cultures remained in the GDP. Again, however, the scaling behaviour indicates that the system was close enough to the phase transition for the critical point to dominate the system behaviour.

There are two caveats that should be considered in the interpretation of our results. First, our analytic results are derived for equilibrium trait distributions, which we likely do not have in our experimental data. However, simulations indicate that the same behaviour can be expected also for transient distributions (Section S3.1). Second, the discarding of outliers, described in Section S2.1, requires careful justification, as it substantially affects the moments of power law distributions. Indeed, the experimental data are subject to measurement errors and both biological and experimental conditions that are not replicated in our theory or simulations. For example, in our theory and simulations cell division is instantaneous, whereas in the experimental data some fraction of the cells are scanned by flow-cytometry during cell division (and these will most likely have large size and high chlorophyll content). This is supported by the fact that a large fraction of flow-cytometry pulses from cells classified as outliers show multimodality, indicating preparation for cell division (see Section S2.1). Additionally, measurement errors occur in the scanning flow cytometer (cells becoming stuck and moving below the expected rate of flow, or data processing errors) that result in anomalously large readings. The small number of high trait value cells, that dominate the higher distribution moments, additionally results in substantial fluctuations particularly in those moments. Cutting off a certain quantile of such “outliers”, however, might also lead to a spurious kink in the moment scaling (similar to the one in Figure 3b) during a transient, even if *α* crosses 1 in a continuous rather than discontinuous manner. A more general type of phase transition, including such continuous changes of *α*, could be defined using a different order parameter (e.g. the fraction of cells with trait *x* > *v*), which then comes with a larger universality class of limiting mechanisms (including, for instance, a hard cut-off preventing cells with trait values *x* < *u* from dividing). Although we see evidence for our ‘discontinuous *α*’ phase transition in our current data, and believe our outlier rejection to be well justified on biological grounds, the more general phase transition can not be definitively ruled out. It would therefore be valuable to focus future experimental work not only on scaling, but also on the mathematical class of limiting processes required to capture growth and division dynamics in unicellular species of interest.

In this manuscript, we have not provided an exhaustive description of the limits of our model class, and focused only on single traits. For more general descriptions, our model class might be enlarged, for example by considering cell *age* as an additional trait to more accurately describe eukaryotic cell division processes. Furthermore, we have focused on cell populations in their exponential growth phase and neglected *cell interactions*, which are salient in natural communities. However, we find the same critical point and phase transition when including a simple type of mean-field “interaction”, that modifies the probability of cell death. As an example of such an interaction, competition for a limited resource might be modeled by subjecting each cell to a death rate that is proportional to some extensive quantity calculated over the whole population (e.g. the total number of cells, or the cumulative value of a certain trait). This kind of interaction would moreover lead to an equilibrium state with a finite number of cells.

Recall that our models are characterised by two relevant parameters, *τ* = (*ω*_2_/*ω*_1_ − 1 and *h* = *u/v*, which encode the natural time- and trait-scales of the trait dynamics. Unveiling the underlying power law exponent *α* from our experiments, through moment scaling, was possible because these parameters could be modified independently. While *τ* appears to be primarily influenced by the nutrients, *h* appears to vary primarily with the light condition (or the species in case of the size distributions). Disentangling the effects of changing *τ* and *h* allowed us to study the scaling of ⟨*x*⟩ with respect to *h*, but our theory also predicts scaling laws with respect to *τ* (see Equations (4) and (5)). In ecological terms, the scaling laws (4) are related to the *resistance* of the system, while the law (5) relates to its *resilience*. Thus *τ* can be used to relate the temporal and trait-scales in the system dynamics. Moreover, the scaling laws in Equations (4) and (5) can be combined to allow model validation without requiring measuring *τ* itself. Such a model validation may be achieved, for example, by measuring the response of trait distributions to external perturbations in an array of chemostats. If the population in each chemostat were kept at a different distance to the critical point (e.g. by careful tuning of nutrient content), the predicted relationship between system responses (e.g. mean and susceptibility) could be observed.

The broader relevance of our work for ecology stems foremost from the dominance of the critical point over the system’s behaviour. This dominance allows system behaviour to be accurately described in terms of universal scaling laws with respect to only two relevant parameters. The universality of the scaling behaviour suggests that it can be expected to occur across diverse systems of growing and dividing individuals. Indeed, scaling relationships, which are evidence of proximity to a critical point, are ubiquitous in ecology, and many might be investigated using the presented approach. We have focused here on chlorophyll-a and light intensity, but many other pairs of trait and environment variables might also be related through our scaling laws, such as size and temperature.

Several well-known ecological phenomena might also be related to our transition. For example, in response to certain (stressful) conditions, many organisms allocate preferentially resources to somatic growth or to reproduction (Heino and Kaitala, 1999; Fischer et al., 2013). Such changes in environmental conditions could change the relative dominance of the rates *ω*_1_ and *ω*_2_, and hence lead to a phase transition with drastic implications for the mean trait. A promising consequence is that if a reliable relationship between an environmental driver and the fundamental rates of our model class could be established for an ecological system, this could be used to predict what would need to happen to either force, reverse, or avoid the transition, with clear applications in environmental management or policy making.

Many scaling relationships are already used directly and indirectly in trait-based modelling studies, particularly in plankton ecology, to constrain or define relationships between traits and the environment, trade-offs, and growth and reproduction of organisms (Litchman and Klausmeier, 2008; Barton et al., 2013; Marañón, 2015; Smith et al., 2011; Merico et al., 2009). It is generally assumed that these scaling relationships are static, with rare exceptions (Merico et al., 2014). Our approach shows quantitatively how some of these scaling relationships might change in dynamic transition regimes of interest, such as environmental resource gradients, and might prove useful in augmenting these existing trait-based modelling approaches.

## Acknowledgements

We are grateful for the continued in-depth discussions with Amos Maritan and Andrea Rinaldo. This work was part of a project supported by SNF (Nr 159660).

