## Supporting Information for "A Universal Phase Transition in Plankton Trait Dynamics"

June 2019

#### Supporting Information Contents

This supplementary material is intended to provide derivations of the theoretical results and more details about the experimental results presented in the main text. Section S1 provides a detailed description of our model class as well as a derivation of the scaling laws associated with its critical point. Section S2 provides more details on the biological experiment where chlorophyll content of phytoplankton was measured in different environments, as well as direct links between the theoretical description and the biological observations. Section S3 contains details concerning our numerical experiments.

#### S1 Phase transition and critical scaling laws

In this section, we investigate the *universal scaling laws* describing the vicinity of the critical point within the mean-field approximation introduced in the main text (c.f. Eq. (1)), and for the class of models that limit low trait values by an increased growth rate for small $x$, and large trait values by an increased division rate for large $x$. We thus set

$$g(x) = \omega_1 x(1 + \tilde{g}(u/x))\,, \quad b(x) = \omega_2(1 + \tilde{b}(x/v))\,, \tag{S1}$$

where $\tilde{g}$ and $\tilde{b}$ are monotonously increasing analytic functions that can be expanded into Taylor series $\tilde{g}(y) = y + \mathcal{O}(y^2)$ and $\tilde{b}(y) = y + \mathcal{O}(y^2)$, around the origin. Parameters $u$ and $v$ define the lower and upper limiting trait values, respectively. The simplest forms of limiting processes are given by the linear functions $\tilde{g}(y) = y$ and $\tilde{b}(y) = y$. We define the two dimensionless parameters $\tau = \omega_2/\omega_1 - 1$ and $h = u/v$, which describe, respectively, the time and trait scales of the theory relative to $\omega_1^{-1}$ and $v$. Henceforth, we work in units where $\omega_1 = 1$ and $v = 1$. Thus, we have a *critical point* at $\tau = h = 0$ (c.f. main text Section 3.1). In the following, we omit the death process for simplicity (i.e. $\omega_3 = 0$ in Eq. (1)), as it does not affect the equilibrium trait distribution we are interested in here.

The critical point marks a *second order phase transition*, where $\langle x\rangle$ plays the role of the *order parameter*. We will show below that the scaling of $\langle x\rangle$ w.r.t. both $|\tau|$ and $h$ can be expressed in the form

$$\langle x\rangle = |\tau|^{\beta} f_{\pm}\left(\frac{h}{|\tau|^{\Delta}}\right)\,, \tag{S2}$$

where the *critical exponents* take the values $\beta = 1$ and $\Delta = 2$. The two different scaling functions $f_{\pm}$, valid for positive and negative $\tau$, respectively, are analytic functions, for small arguments, and both $f_+(y)/y$ as well as $f_-(y)$ converge to positive constants when $y \to 0$. We shall also see that this phase transition is a consequence of a *jump* of exponent $\alpha$ in Eq. (2) at the transition point $\tau = 0$.

Upon integrating Eq. (1), we find that, once the population has reached a steady exponential growth, its growth rate is given by

$$\omega = \langle b(x) \rangle \,, \tag{S3}$$

where the brackets denote the expectation value with respect to the normalised equilibrium trait distribution. Multiplying Eq. (1) first with $x$ before integration yields an equation for the dynamics of the moments,

$$\frac{d}{dt}\langle x \rangle_t = \langle g(x) \rangle_t - \langle x \rangle_t \langle b(x) \rangle_t \,. \tag{S4}$$

Plugging Eqs. (S1) into Eq. (S4) yields, for the equilibrium distribution,

$$\langle x \tilde{g}(h/x) \rangle = \langle x \rangle (\tau + \langle \tilde{b}(x) \rangle) \,. \tag{S5}$$

Let us first consider the GDP ($\tau < 0$). Due to the assumptions made above about $\tilde{g}(y)$, the l.h.s. of Eq. (S5) scales like $h$, for small $h$. Hence, in the limit $h \to 0$, $\langle \tilde{b}(x) \rangle \to |\tau|$. The case $\langle x \rangle \to 0$ would entail $\langle \tilde{b}(x) \rangle \to 0$ and thus contradict positivity of the l.h.s. of Eq. (S5). Hence, we find that $\langle x \rangle \sim h^0$. For the scaling of the higher moments, we need to derive exponent $\alpha$ in Eq. (2). This exponent is derived from Eq. (1), plugging in a pure power-law $\psi(x,t) = x^{-\alpha}$ and letting $u \to 0$ and $v \to \infty$. We then find that $\alpha$ solves

$$\omega + \omega_1(1-\alpha) + \omega_2(1 - 2^{2-\alpha}) = 0 \,. \tag{S6}$$

Using Eq. (S3), Eq. (S1), and the fact that $\langle \tilde{b}(x) \rangle \to -\tau$, we conclude that the population growth rate satisfies $\omega = 1 + \mathcal{O}(\tau^2)$. Plugging this into Eq. (S6) and keeping only terms up to linear order in $\tau$ we find that

$$\alpha - 2 = (1+\tau)(1 - 2^{-(\alpha-2)}) \,. \tag{S7}$$

This equation has two solutions, for $\tau \lesssim 0$ (i.e. $\tau < 0$ and $\tau \approx 0$), namely $\alpha = 2$ and another one that is close to but smaller than 1. The first solution would lead to the contradiction $\langle x \rangle \to 0$, which means that $\alpha \lesssim 1$. Thus we find that, for $h \to 0$,

$$\langle x^k \rangle = \frac{\int_0^\infty x^{k-\alpha} \mathcal{F}(h/x, x)}{\int_0^\infty x^{-\alpha} \mathcal{F}(h/x, x)} \propto 1 - \alpha \,. \tag{S8}$$

From Eq. (S7) we deduce that, near criticality, $\alpha = 1 + |\tau|/(1 - \ln 2)$. Thus

$$\langle x^k \rangle \sim |\tau| \,, \tag{S9}$$

for all $k \geq 1$. This also implies that $\beta = 1$ in Eq. (S2).

In the DDP ($\tau > 0$), on the other hand, we derive from Eq. (S5) that $\langle x \rangle \sim h/\tau$, for $h \ll \tau$. Thus, the second critical exponent defined in Eq. (S2) is $\Delta = 2$. From Eq. (S3) and Eq. (S1) we further derive $\omega = 1 + \tau$, when $h \to 0$. Thus, from Eq. (S6) we find that $\alpha$ satisfies

$$\alpha - 1 = 2(1+\tau)(1 - 2^{-(\alpha-1)}) \,. \tag{S10}$$

This equation has two solutions, namely $\alpha = 1$ and another one that is close to but larger than 2. The first solution is incompatible with $\langle x \rangle = \mathcal{O}(h)$, which means that $\alpha \gtrsim 2$. *At the transition point, $\alpha$ thus jumps from 1 to 2.* For the higher moments we find, using Eq. (2), *anomalous scaling* with respect to $h$:

$$\langle x^k \rangle \sim \begin{cases} h^k\,, & k < \alpha - 1 \\ h^{\alpha-1}\,, & k \geq \alpha - 1\,. \end{cases} \tag{S11}$$

The simplest member of our model class has $\tilde{g}(y) = y$ and $\tilde{b}(y) = y$. In this case, $\langle x \rangle$ can be calculated exactly:

$$\langle x \rangle = \frac{-\tau + \sqrt{\tau^2 + 4h(\tau+1)}}{2(\tau+1)}\,, \tag{S12}$$

and the scaling laws Eq. (S2) and critical exponents $\beta = 1$ and $\Delta = 2$ can be verified immediately.

From Eq. (S4) we further derive the dynamics of fluctuations around the stationary solution. Setting $\langle x \rangle_t = \langle x \rangle + \epsilon(t)$, we find that, for $h \ll |\tau|$,

$$\dot{\epsilon}(t) = -|\tau|\epsilon(t) + \mathcal{O}(|\tau|) + \mathcal{O}(\epsilon(t)^2)\,. \tag{S13}$$

This implies that perturbations decay exponentially with time scale $|\tau|$ (or $|\omega_1 - \omega_2|$). At criticality, where this time scale vanishes, the return to equilibrium follows a much slower algebraic decay $\epsilon(t) \sim t^{-1}$ (*critical slowing down*).

### S2 Adaptation of chlorophyll content in different light intensities

#### S2.1 Evidence for phase transition in all cultures

Here we show evidence that the phase transition occurs not only in the directly illuminated *Pseudokirchneriella* cultures that were shown in the main text, but also in the *Microcystis* cultures, and in shaded cultures of both species (which received only light filtered by the competitor).

Initially, all cultures appear to be in the GDP, where we observe fast exponential population growth (Figure S1) and trivial moment scaling of the form $\langle x^k \rangle \sim \langle x \rangle^k$, for all observable positive moments (Figures S2, S3). Later in the experiment, all cultures appear to transition to the DDP, where the population growth rate decreases, and the anomalous moment scaling $\langle x^k \rangle \sim \langle x \rangle^{\gamma k}$, with $\gamma < 1$, is observed. In all cultures, we observe a strong decrease in mean chlorophyll content throughout the experiment (Figure S4).

It is evident that the features characteristic for our phase transition (change in population growth rate and moment scaling) can already be observed before the systems reach an equilibrium trait distribution. In Section S3.1, we will show through numerical simulation that this is also the case for our model.

Here, we show all data including also the lowest light intensity cultures that were not shown in the main text. The cultures from this light intensity all show very slow population growth (see Figure S1) as well as little change in the mean chlorophyll-a content over time (see Figure S4). From population and trait dynamics it is therefore unclear whether

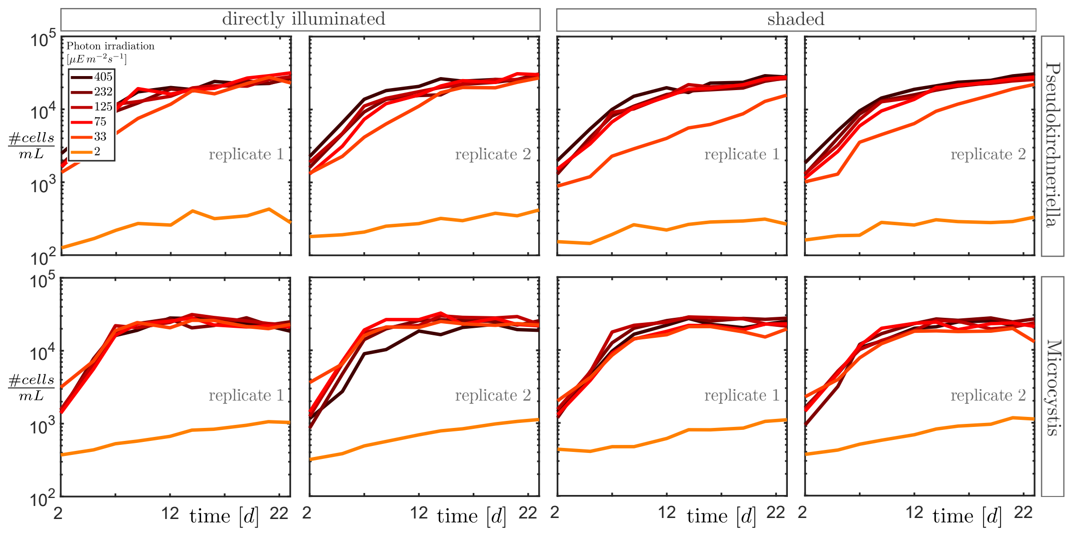


Figure S1: Population growth in all directly illuminated and shaded cultures of both cell types. All cultures clearly show the transition from fast to slow population growth.

these cultures underwent the phase transition. Nevertheless, we present the data here for completeness and show that the moment scaling is robust to the inclusion of these cultures.

Before the computation of the moment scaling shown in Figures 3, S2, and S3, outliers (between $\approx 1\%$ and $\approx 5\%$ cells with largest chlorophyll content) were removed from the data to reduce the noise and reveal the moment scaling (the moment scaling before the removal of outliers is shown in Figures S5 and S6). These outliers dominate particularly the higher moments and obscure the moment scaling completely. They are thought to originate mainly from cells in the process of cell division at the time of measurement, as well as faulty measurements (see also the discussion in Section 4). Indeed, we find evidence in the flow-cytometry signals of the cells classified as outliers that a large fraction of these cells are in the process of division. Between $\approx 30\%$ and $\approx 50\%$ of the flow-cytometry pulses (recording the forward- and sideward scattered light and chlorophyll fluorescence signals through time) within the outliers show multimodality, indicating that the cell content is being split up in preparation for division. Within the same number of cells below the threshold (above which cells are classified as outliers), only between $\approx 5\%$ and $\approx 20\%$ of the flow-cytometry pulses show multimodality. The outliers are classified as the data points that are above the quantile $q = (q_u - q_l) \cdot 1.5 + q_u$, where $q_l$ is the 25% quantile and $q_u$ is the 75% quantile. The percentage of outliers is typically larger in the GDP and lower in the DDP. Such a removal of outliers could potentially change the moment scaling. If a constant fraction of outliers above a quantile $q_u$ is removed, in the equilibrium trait distributions of our model, the moment scaling would also in the DDP revert to a trivial moment scaling. This is because any upper quantile will scale as $q_u \sim u$, dominating the scaling of all higher moments. However, it appears that during the transients, a difference in scaling between the first and the higher moments can still be observed, even after removing the outliers. An explanation for this may be that after the transition, a significant amount of cells is pushed towards lower trait values, so that the mean $\langle x \rangle$ begins to scale with $u$, while enough cells are still large enough for the upper quantile (and thus the higher moments) to be dominated by the

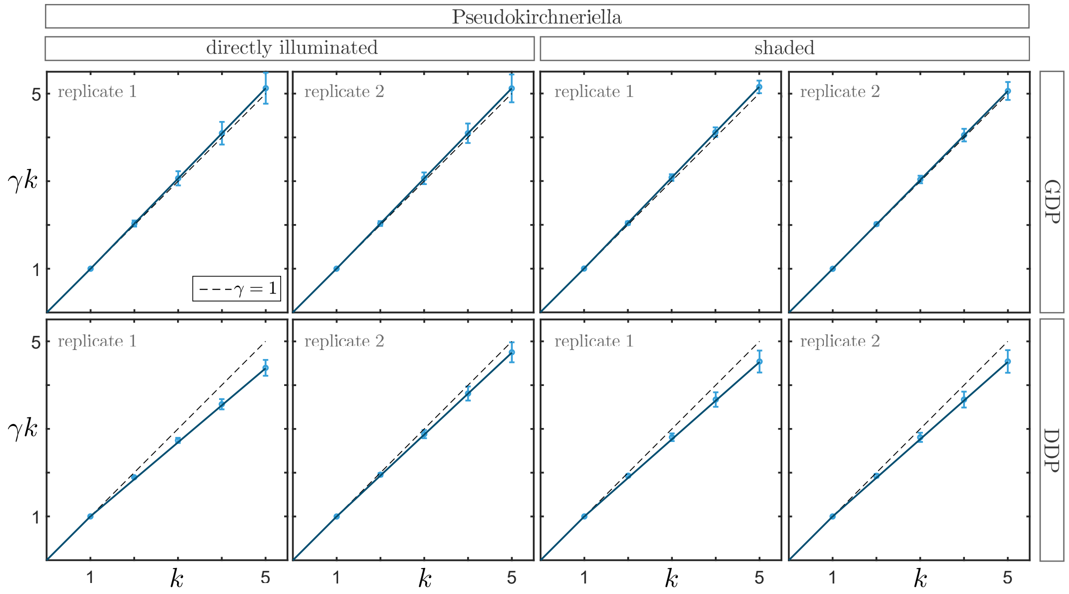


Figure S2: Moment scaling in all directly illuminated and shaded *Pseudokirchneriella* cultures. Plots were obtained in the same way as Figure 3 in the main text.

$v$-scale. The moment scaling in the GDP should not be affected, since all moments, as well as any quantile, will scale with $v$.

The uncertainty in the moment scaling in the GDP remains relatively large in some cultures even after the removal of outliers. This may be due to the fact that the cells are still adapting to the experimental environment (individual cells can influence the higher moments strongly, and close to the critical point, convergence may take particularly long, see Section S1). However, the trend of close to trivial moment scaling initially, and anomalous moment scaling later, can be seen in all cultures. The trivial moment scaling in the GDP is evidence that $\alpha$ stays close to 1 throughout the initial phase of the experiment (see Section S2.2). In the DDP, the change in moment scaling occurs for moments $k > \alpha - 1 \approx 1$, which is evidence that $\alpha$ stays close to 2 (see Eqs. (S11), (S14)).

In Section S2.2 we show how both the change in population growth and the change in moment scaling with respect to the first moment follow directly from our theoretical results. The fact that the phenomenology of the phase transition persists across these different species and environmental conditions is further evidence for the universality of this transition.

#### S2.2 Link to theoretical results

The change in population growth rate and moment scaling exhibited by all cultures are evidence for the phase transition described above. As shown in Section S1, for the model without a death process, the population growth rate is given by $\omega = 1 + \mathcal{O}(\tau^2)$ in the GDP, and $\omega = 1 + \tau$ in the DDP, up to corrections of order $h$. In terms of the original frequency parameters, this becomes $\omega \approx \omega_1$ in the GDP and $\omega \approx \omega_2$ in the DDP. Close to the critical point, the steady-state population growth rate can thus be expressed as $\omega \approx \max(\omega_1, \omega_2)$.

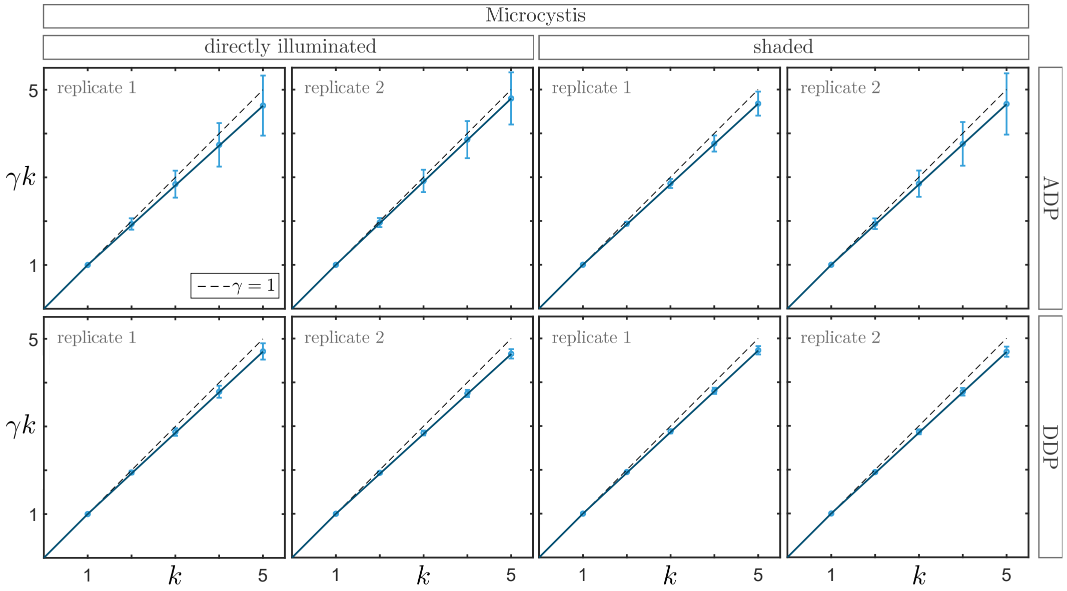


Figure S3: Moment scaling in all directly illuminated and shaded *Microcystis* cultures. Plots were obtained in the same way as Figure 3 in the main text.

This is because when cells grow (i.e. increase their traits, such as chlorophyll content) fast, they will quickly reach the limiting $v$-scale and divide, so their somatic growth rate will dominate the population growth rate. On the other hand, when the somatic growth rate is lower, the uniform (trait-independent) division rate will be dominant. Under the assumption that $\omega_2$ stays smaller than the initial $\omega_1$ of the GDP, this results in a decrease in population growth rate after the transition. If a uniform death rate $\omega_3$ (c.f. Eq. (1)) is included, the population growth rate is reduced to $\omega \approx \omega_1 - \omega_3$ in the GDP and $\omega \approx \omega_2 - \omega_3$ in the DDP. If $\omega_3$ is larger in the DDP than in the GDP, and $\omega_3$ stays roughly the same or is smaller in the DDP than in the GDP, this will increase the difference between population growth rates in the GDP and the DDP.

The change in moment scaling with respect to the control parameter $h$ directly translates to our observed scaling with the first moment. In terms of the original trait parameters of our model, Eq. (S11) translates to

$$\langle x^k \rangle \sim \begin{cases} u^k\,, & k < \alpha - 1\,, \\ v^{k-(\alpha-1)} u^{\alpha-1}\,, & k \geq \alpha - 1\,. \end{cases} \tag{S14}$$

In order to achieve the observed patterns, a scaling relationship of both $u$ and $v$ with the relevant resource, in our case the light intensity $\ell$, has to be assumed (that is, $u \sim \ell^{\mu}$ and $v \sim \ell^{\nu}$). This expresses the intuition that the light availability determines the 'optimal' trait scales in both phases.

It follows that in the GDP close to the critical point, where $\alpha \lesssim 1$, the mean scales like $\langle x \rangle \sim \ell^{\nu+(\alpha-1)(\mu-\nu)}$. The scaling of the higher moments with $\langle x \rangle$ is thus calculated as

$$\langle x^k \rangle \sim \ell^{\nu k+(\alpha-1)(\mu-\nu)} \sim \langle x \rangle^{(\nu k+(\alpha-1)(\mu-\nu))/(\nu+(\alpha-1)(\mu-\nu))}\,.$$

Since $\alpha - 1 \approx 0$, this results, approximately, in the trivial moment scaling $\langle x^k \rangle \sim \langle x \rangle^k$.

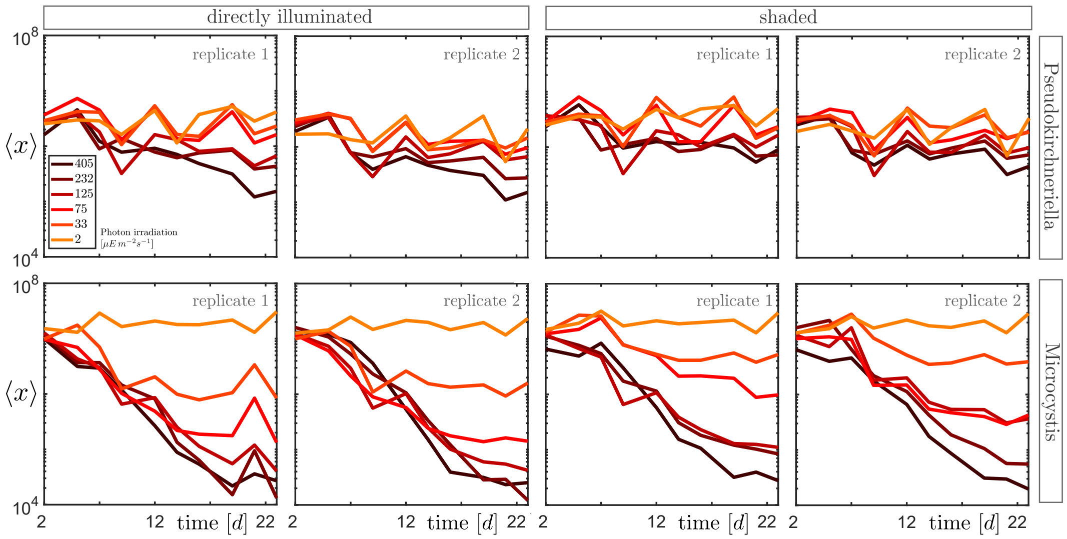


Figure S4: Mean chlorophyll content development in all directly illuminated and shaded cultures of both cell types. All cultures show a strong decrease in mean trait.

In the DDP, where $\alpha \gtrsim 2$, the mean scales with $u$ as $\langle x \rangle \sim u \sim \ell^{\mu}$, while the higher moments still scale like

$$\langle x^k \rangle \sim \ell^{\nu k+(\alpha-1)(\mu-\nu)} \sim \langle x \rangle^{(\nu/\mu)k+(\alpha-1)(1-\nu/\mu)},$$

for $k > 1$. This becomes, for $\alpha \gtrsim 2$, $\langle x^k \rangle \sim \langle x \rangle^{1+(\nu/\mu)(k-1)}$ which results in the observed anomalous moment scaling with $\gamma = \nu/\mu$.

### S3 Numerical experiments

#### S3.1 Simulations of the phase transition

In order to corroborate our theoretical results, we ran numerical experiments with a model from our universality class (where the growth and division rates are in the class defined in Eq. (S1)). We used the simplest of those models, where the growth, division, and death rates are given by, respectively, $g(x) = \omega_1(u + x)$, $b(x) = \omega_2(1 + x/v)$, and $d(x) = \omega_3$. Our data exhibit a strong decrease in the mean chlorophyll content (c.f. Figure S4), while the population growth rate is relatively low, especially after the transition (c.f. Figure S1). A non-zero death rate was thus needed to simultaneously reproduce trait and population dynamics. Using our adapted Gillespie algorithm (c.f. Section S3.2), experimental data as initial distributions, and parameter values consistent with our theoretical requirements (discussed below), the phase transition and associated scaling behaviour could be reproduced consistently. The aim of these simulations was not a quantitative reproduction of a specific data set, but qualitative reproduction of the phenomenology of the phase transition. Additionally, we find that even in the transients, where the distributions have not converged to a steady state yet, the transition in population growth rate and moment scaling can already be observed, as is the case in our experimental data.

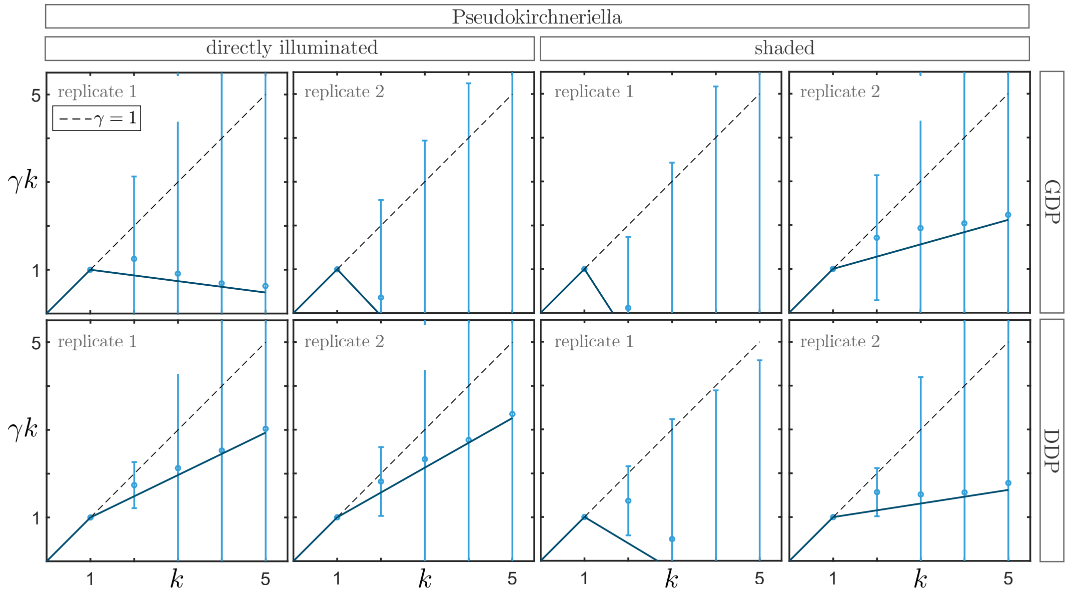


Figure S5: Moment scaling in all directly illuminated and shaded *Pseudokirchneriella* cultures, *before* the removal of outliers. Moment scaling is completely obscured by the inclusion of outliers, so that the phase transition becomes unrecognisable. Outliers are thought to originate from cells in the process of division and inaccurate measurements (see Section 4).

Both the characteristic transition in population growth rate (Figure S7a) and the decrease in the mean chlorophyll content (Figure S7b) are reproduced. The transition in moment scaling (Figure S8), resulting from the loss of single-scale dominance and data collapse (Figure S9), are also observed. For the computation of the moment scaling we did not remove any outliers here, since the biological and technical origins of these outliers (see Section 4) are not present in our simulations.

The simulations were run with parameter values that are consistent with our theoretical requirements. We require that $\tau < 0$ and $|\tau| \approx 0$ in the GDP (first part of the experiment), in order to obtain $\alpha \approx 1$ and trivial moment scaling (c.f. Section S2.2). In the DDP (second part of the experiment), we require $\tau > 0$ and $|\tau| \approx 0$, in order to obtain $\alpha \approx 2$ (c.f. Section S2.2). The parameter $\tau$ also needs to be equal for all cultures, and a scaling relationship of both $u$ and $v$ with the relevant resource, the light, needs to exist (c.f. Section S2.2).

In the GDP, starting from the distributions measured in the experiment, the self-similarity and trivial moment scaling is robustly (i.e. mostly independent of the specific rate parameter values) preserved in the transient distributions of the model. The transition from GDP to DDP is initiated when a certain particle number is reached, upon which all rates are changed instantaneously. In the DDP, the change in moment scaling, due to the emergence of the second relevant trait scale $u$, is observed very shortly after the phase transition.

The exact parameter values used here are shown in Table S1. The rates $\omega_1$ and $\omega_2$ were inferred mainly from the trait dynamics (speed of moment evolution), while $\omega_3$ was inferred from the other two rates together with the population growth rate (c.f. Section S2.2). The trait scales $u$ and $v$ were chosen to reproduce roughly the experimentally observed decrease

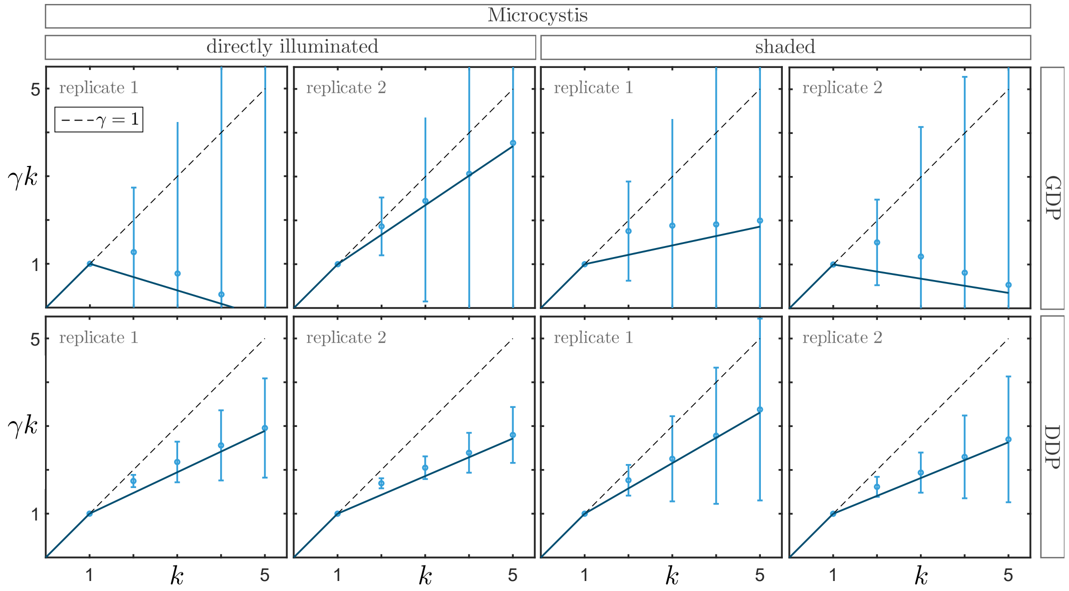


Figure S6: Moment scaling in all directly illuminated and shaded *Microcystis* cultures, *before* the removal of outliers. Moment scaling is completely obscured by the inclusion of outliers, so that the phase transition becomes unrecognisable. Outliers are thought to originate from cells in the process of division and inaccurate measurements (see Section 4).

in mean chlorophyll content and the experimentally observed moment scaling exponent $\gamma k$ (c.f. Section S2.2). In both phases, the parameter values correspond to a system close to the critical point. The rate values are very similar in both phases, only $\omega_1$ decreased and $\omega_3$ increased in the DDP, corresponding to slower somatic growth and higher death rate. Both these changes appear to be biologically plausible and consistent with experiments. The rate parameters here are set equal for all cultures, but could be varied across the cultures. We only need to ensure that all cultures have roughly the same $\tau$ and $\omega$, and that the rates are large enough for a significant number of cells to come under the influence of $u$ in the DDP.

While the scaling behaviour is reproduced well by the model, there are still differences between this specific model and the experimental measurements, for instance the exact trait dynamics (Figure S7) and the shape of the trait distributions (Figure S9). This is not surprising, given that these simple growth and division mechanisms are likely not a perfect description of the complex behaviour of these cells. Additionally, in our simulations the transition from GDP to DDP happens instantaneously, while in reality, the rates would naturally change continuously in time. While the details of the models influence details of the trait distributions and the behaviour during the transients, they do not influence the observed phase transition and the scaling behaviour around the critical point.

#### S3.2 Generalised Gillespie algorithm for many particle systems with variable rates

Simulating such many particle systems can be helpful or even necessary, for instance in cases where trait distribution solutions cannot be found analytically. The Gillespie algorithm

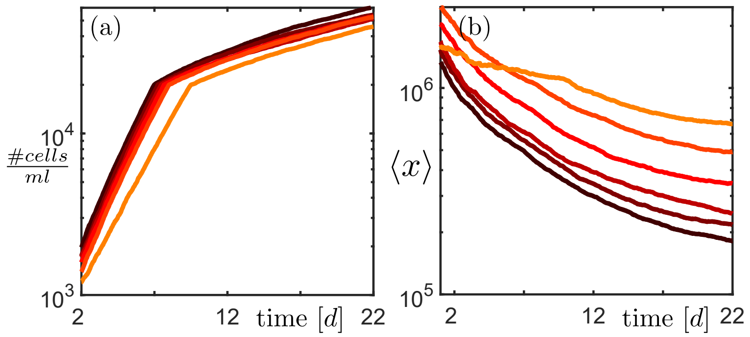


Figure S7: Simulation results for six cultures (different colors), where the phase transition (i.e. the parameter switch) is initiated when a certain population size is reached. (a) Time development of population size. (b) Time development of first moment.

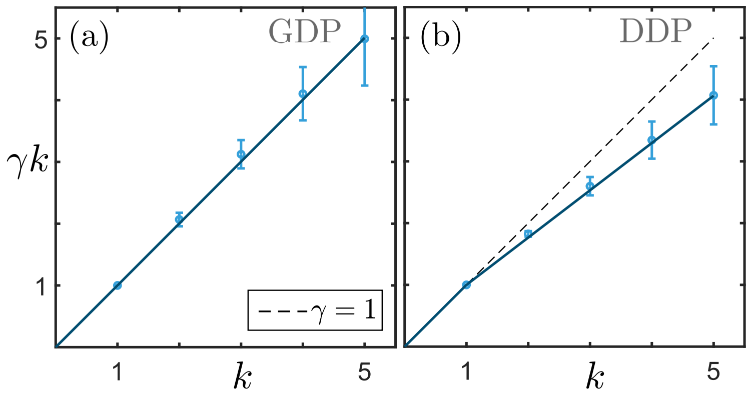


Figure S8: Moment scaling during the simulation shown in Figure S7, (a) before the transition (GDP), (b) after the transition (DDP).

(Gillespie, 1976) offers an efficient way to simulate large numbers of particles with constant reaction rates. Rather than simulating forward in time with constant time increments, in Gillespie's algorithm the distribution of reaction event times is computed, and event times are then drawn from this distribution. If the reaction rates are constant, the probability for an event to happen grows exponentially in time.

In our models, the only 'reactions' are cell division and cell death events. From the division rate $b(x(t))$ and the death rate $d(x(t))$, we can compute the 'survival probability' $s(t)$, which is the probability for the cell to survive (i.e. not divide and not die) until time $t$. Since these rates are in general not constant but dependent on the state of the cell, the trait $x(t)$, the survival probability is no longer an exponential function of time as in the original Gillespie algorithm. However, for certain cases of $g$ and $b$, the survival probability can still be computed analytically.

We show the computation here, as an example, for the simplest model from our model class (S1), where the growth and division rates are given by, respectively,

$$g(x) = \omega_1 (u + x) \,, \tag{S15}$$

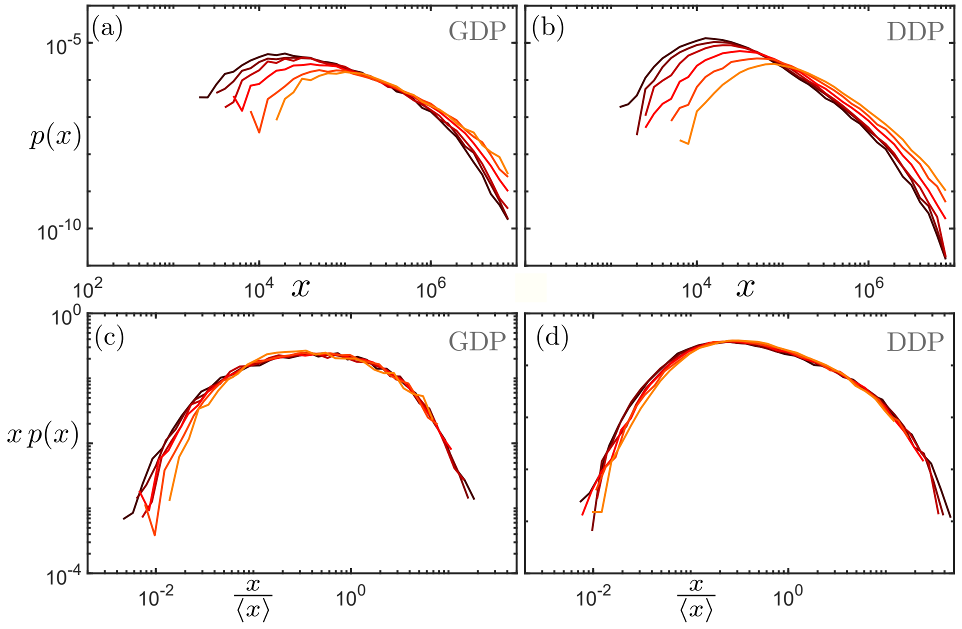


Figure S9: Simulated trait distributions before and after the transition. (a) Distributions before the transition (GDP). (b) Distributions after the transition (DDP). (c) Rescaled distributions in the GDP. (d) Rescaled distributions in the DDP.

$$b(x) = \omega_2(1 + x/v)\,. \tag{S16}$$

Looking first at a single process, cell division, the survival probability distribution $s(t)$ is defined through the division rate as

$$\frac{\mathrm{d}s(t)}{\mathrm{d}t} = -b(x(t))s(t)\,. \tag{S17}$$

Since the cells grow according to Eq. (S15), the trait $x(t)$ of a single cell at time $t$ is given by

$$x(t) = (x_0 + u)e^{\omega_1 t} - u\,, \tag{S18}$$

where $x_0$ is the trait value of the cell at $t = 0$. Eq. (S17) then becomes

$$\frac{\mathrm{d}s(t)}{\mathrm{d}t} = -\omega_2\left(1 + \tfrac{x_0+u}{v}e^{\omega_1 t} - \tfrac{u}{v}\right)s(t)\,, \tag{S19}$$

which can be solved to give the survival probability distribution

$$\log s(t) = \frac{\omega_2}{\omega_1}\frac{x_0 + u}{v} + \omega_2\left(\frac{u}{v} - 1\right)t - \frac{\omega_2}{\omega_1}\frac{x_0 + u}{v}e^{\omega_1 t}\,. \tag{S20}$$

To generate division times from this distribution, one can now draw a random value $k$ from a uniform distribution in the interval $[0, 1]$ (i.e., 'draw a survival probability') and invert Eq. (S20) to obtain the corresponding division time $t_b$. That is, one solves

Table S1: Parameter values in model simulations.

| Phase | $\omega_1$ | $\omega_2$ | $\tau$ | $\alpha$ | $\omega_3$ |
|---|---|---|---|---|---|
| GDP (all cultures) | 0.71 | 0.7 | $-0.02$ | $\approx 0.95$ | 0.41 |
| DDP (all cultures) | 0.67 | 0.7 | 0.05 | $\approx 2.16$ | 0.7 |

| Culture | $u$ | $v$ |
|---|---|---|
| 1 | $1.34 \cdot 10^4$ | $2.65 \cdot 10^6$ |
| 2 | $1.65 \cdot 10^4$ | $3.06 \cdot 10^6$ |
| 3 | $2.03 \cdot 10^4$ | $3.54 \cdot 10^6$ |
| 4 | $3.08 \cdot 10^4$ | $4.73 \cdot 10^6$ |
| 5 | $4.66 \cdot 10^4$ | $6.33 \cdot 10^6$ |
| 6 | $7.05 \cdot 10^4$ | $8.46 \cdot 10^6$ |

$$\log s(t_b) = \log k \tag{S21}$$

for $t_b$. For general $g$ and $b$, this may be hard to compute. However, in systems with many cells the relevant division times will generally be very small (e.g., for the rates chosen in Section S3.1 they are $\approx 10^{-4}$), so we can expand the exponential in Eq. (S20) in $t$ up to second order and solve the resulting equation

$$\omega_1\omega_2 \frac{(x_0+u)}{2v} t_b^2 + \omega_2 \left(\frac{x_0}{v}+1\right) t_b + \log k = 0\,, \tag{S22}$$

from which the division time is then given by

$$t_b = -\frac{1}{\omega_1}\frac{x_0+v}{x_0+u} + \frac{1}{\omega_1}\frac{v}{x_0+u}\sqrt{\left(\frac{x_0}{v}+1\right)^2 - 2\frac{\omega_1}{\omega_2}\frac{x_0+u}{v}\log k}\,. \tag{S23}$$

This expression is computed for all cells, and the smallest division time determines the next division event.

Additional processes are treated the same way, by computing the survival probabilities of all cells under these processes, and then selecting out of all events the event with the smallest event time. For events that occur with the same probability for all cells, such as a uniform death rate $d(x) = \omega_3$, the survival probability decays exponentially,

$$s(t) = e^{-\omega_3 t}\,. \tag{S24}$$

Drawing a random value $k$ and solving Eq. (S21) for all cells individually, however, can be computationally expensive for large cell numbers. Instead we can use the following property of the exponential distribution to speed up the simulation. For $N$ independent exponentially distributed random variables with rate parameters $\{\omega_3(1), ..., \omega_3(N)\}$, the *minimum* of these random variables is also exponentially distributed, with rate parameter $\omega_N = \sum_{i=1}^{N} \omega_3(i)$. The minimum survival time of all $N$ cells with the constant rate $\omega_3$ is thus given by

$$s(t) = e^{-\omega_3 N t}\,, \tag{S25}$$

which has to be evaluated only once to compute the minimum survival time, upon which a cell can be chosen (with uniform probability) to experience the event accordingly.
